# Bone-targeted lncRNA OGRU alleviates unloading-induced bone loss via miR-320-3p/Hoxa10 axis

**DOI:** 10.1101/745430

**Authors:** Ke Wang, Yixuan Wang, Zebing Hu, Lijun Zhang, Gaozhi Li, Lei Dang, Yingjun Tan, Xinsheng Cao, Fei Shi, Shu Zhang, Ge Zhang

## Abstract

Although the underlying molecular mechanism of unloading-induced bone loss has been broadly elucidated, the pathophysiological role of long noncoding RNAs (lncRNAs) in this process is unknown. Here, we identified a novel lncRNA, OGRU, a 1816-nucleotide transcript with significantly decreased levels in bone specimens from hindlimb-unloaded mice and in MC3T3-E1 cells under clinorotation unloading conditions. OGRU overexpression promoted osteoblast activity and matrix mineralization under normal loading conditions and attenuated the suppression of MC3T3-E1 cell differentiation induced by clinorotation unloading. Furthermore, this study found that supplementation of pcDNA3.1(+)-OGRU via (DSS)_6_-liposome delivery to the bone formation surfaces of hindlimb-unloaded (HLU) mice partially alleviated unloading-induced bone loss. Mechanistic investigations demonstrated that OGRU can function as a competing endogenous RNA (ceRNA) to facilitate the protein expression of Hoxa10 by competitively binding miR-320-3p and subsequently promote osteoblast differentiation and bone formation. Taken together, the results of our study provide the first clarification of the role of the OGRU in unloading-induced bone loss through the miR-320-3p/Hoxa10 axis, suggesting an efficient anabolic strategy for osteoporosis treatment.

## Introduction

Bone remodeling is a complex process orchestrated by the dynamic balance between osteoblast-mediated bone formation and osteoclast-mediated bone resorption.^1^ This balance can be regulated by a variety of factors, such as hormones, cytokines and mechanical stimuli.^2^ The effect of mechanical stimuli on bone remodeling has been described by Wolff’s law: the structure and function of bone are adapted to its mechanical environment.^3^ Accumulating evidence has shown that prolonged mechanical unloading by long-duration space flight and extended bed rest can induce disuse osteoporosis in humans.^4, 5^ The most devastating consequence of osteoporosis is fractures, which are associated with an enormous burden of health care costs, morbidity and mortality. Existing antiosteoporotic drugs can effectively reduce the incidence of fractures in patients with osteoporosis. However, major side effects of these drugs, such as osteonecrosis, hypercalcemia and thromboembolic diseases, can be extremely harmful to human health. Therefore, it is necessary to further explore the molecular mechanism of bone loss to develop safer and more effective antiosteoporotic drugs.^6^

HLU mice, an established model used to simulate skeletal unloading, show irreversible bone loss via the inhibition of bone formation and the promotion of bone resorption.^5, 7–9^ In vitro experiments in ground-based facilities for simulating microgravity unloading, such as 2D clinostat, random positioning machine (RPM), rotating wall vessel (RWV) bioreactor and magnetic levitator, indicated that the dysfunction of osteoblasts caused by unloading can disturb their ability to translate mechanical loading into biochemical signals, which can partly explain the occurrence of unloading-induced bone loss,^10–13^ and many intracellular signaling pathways have been reported to participate in this complex process.^5, 8, 14^ To further understand the pathophysiology of unloading-induced bone loss and identify new biological targets, we selected 2D clinorotation and HLU models for simulating unloading conditions in vitro and in vivo.

Whole-genome sequencing results show that approximately 93% of the sequences in the human genome can be transcribed, among which no more than 2% have protein-coding ability; 98% of the transcriptome consists of noncoding RNAs without protein-coding potential.^15^ Long noncoding RNAs are transcripts of more than 200 nucleotides in length, without long open reading frames but often with mRNA structural features (a 5’ cap and a polyA tail).^16^ Previous studies have found that lncRNAs can participate in a wide range of biological processes by regulating gene expression at the transcriptional, posttranscriptional and epigenetic levels.^17^ For example, lncRNAs can induce a repressive chromatin state by recruiting chromatin-modifying complexes, such as PRC2, Mll, PcG and G9a methyltransferase, to specific genomic loci.^18–22^ Furthermore, lncRNAs can recruit and modulate the activity of the RNA-binding protein TLS and subsequently cause the repression of cyclin D1 transcription in human cell lines.^23^ Moreover, many cytoplasmic lncRNAs can competitively bind to microRNAs by functioning as competitive endogenous RNAs (ceRNAs) and thereby regulate protein translation.^24–27^ Importantly, although the roles of lncRNAs in the process of osteogenesis have been widely reported,^28–30^ only a few studies have shown that lncRNAs are involved in the occurrence and development of unloading-induced bone loss.^31, 32^

In the present study, we found that lncRNA OGRU is downregulated during unloading and upregulated during osteoblast differentiation. OGRU overexpression promoted osteoblast activity and matrix mineralization under normal loading and attenuated the suppression of MC3T3-E1 cell differentiation by unloading conditions in vitro. In an in vivo experiment, a (DSS)_6_-liposome delivery system was used to deliver pcDNA3.1(+)-OGRU specifically to bone formation surfaces.^33, 34^ The results showed that supplementation with (DSS)_6_-liposome-OGRU can effectively promote osteoblastic bone formation, increase bone mass, and improve bone microarchitecture and biomechanical properties in HLU mice. In addition, OGRU facilitated the protein expression of Hoxa10 by competitively binding miR-320-3p, which in turn promoted osteoblast differentiation and bone formation. In summary, our study identified a new regulator of unloading-induced bone loss and may establish a potential therapeutic strategy against osteoporosis.

## Results

### OGRU is a novel lncRNA sensitive to unloading and is upregulated during osteoblast differentiation of MC3T3-E1 cells

To identify unloading-sensitive lncRNAs, a microarray assay was performed in our previous work to detect differential expression of lncRNAs in MC3T3-E1 cells under clinorotation unloading condition for 48 h.^35^ The expression of candidate lncRNAs selected from the microarray assay data according to the fold change in expression was further validated by qRT-PCR. The results showed that NONMMUT068562 (named OGRU) was markedly decreased in the clinorotation unloading group and that this change persisted for at least 72 h in MC3T3-E1 cells (Figure 1A, B). In addition, OGRU was also markedly downregulated in the tibias of HLU mice (Figure 1C). To identify whether OGRU is potentially involved in osteoblast differentiation, MC3T3-E1 cells were cultured in osteogenic medium for 21 d. We found that the expression of OGRU increased gradually and was positively associated with the mRNA and protein expression of the osteogenic marker genes Osx and Runx2 during osteoblast differentiation (Figure 1D, E). Thus, this uncharacterized lncRNA was selected for further study. Next, 5’ and 3’ rapid amplification of cDNA ends (RACE) assays showed that OGRU was a 1816-nucleotide transcript comprising two exons and located on chromosome 9, and northern blot analysis further confirmed the expected size (Figure S1A, B). OGRU displayed no protein-coding potential, as predicted by the Coding Potential Calculator (CPC) and the Coding-Potential Assessment Tool (CPAT) (Table S4). In summary, we identified a novel lncRNA, OGRU, which exhibited significantly decreased expression during unloading both in vitro and in vivo and was associated with osteoblast differentiation of MC3T3-E1 cells.

**Figure 1.**
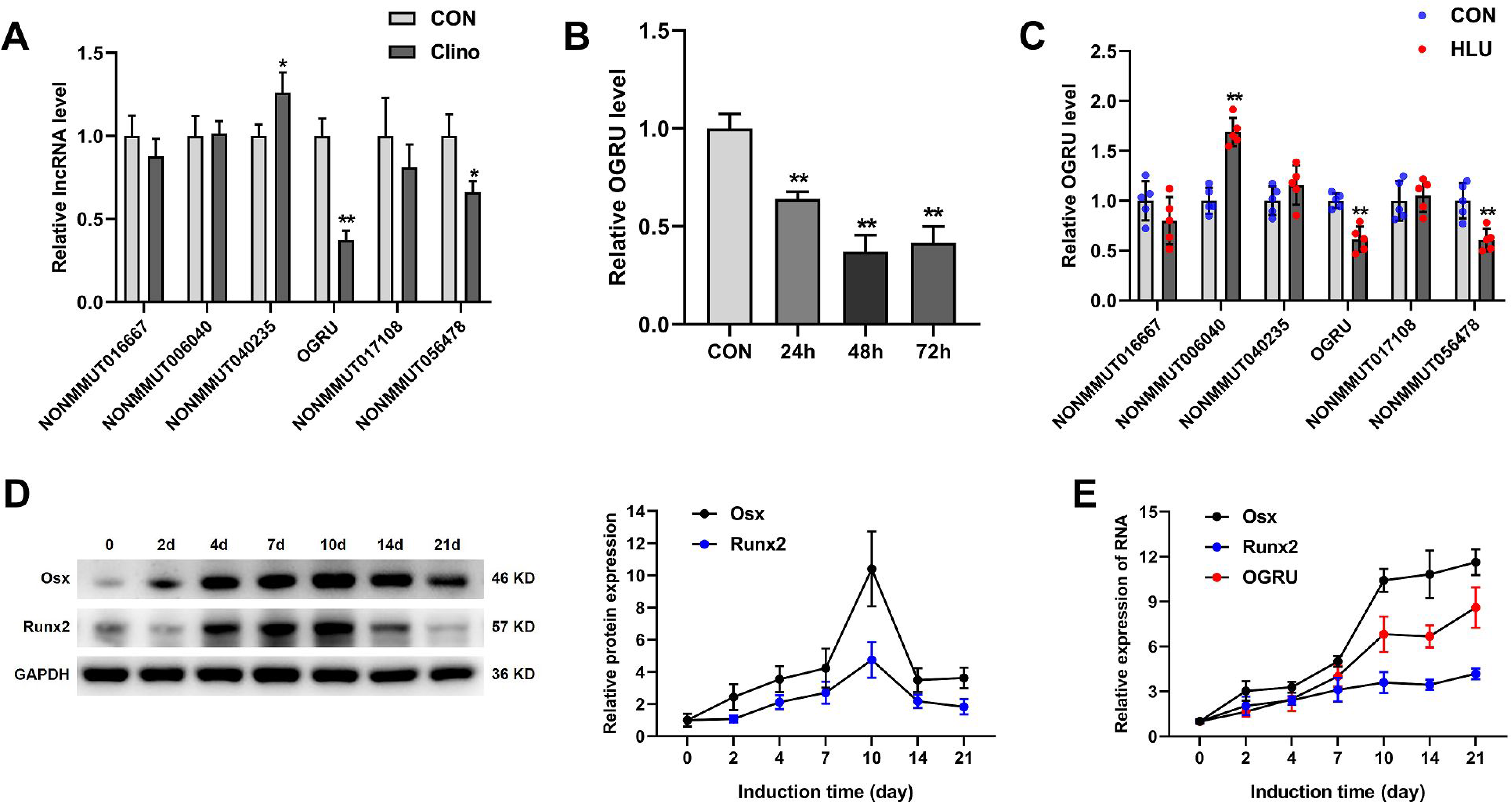
OGRU is sensitive to unloading and is upregulated during osteoblast differentiation of MC3T3-E1 cells. **(A)** Differential expression of lncRNAs selected from previous microarray assay data was determined by qRT-PCR in MC3T3-E1 cells under clinorotation unloading condition for 48 h (n=3). **(B)** qRT-PCR analysis of OGRU expression in MC3T3-E1 cells under clinorotation unloading condition for 24, 48 and 72 h (n=3). **(C)** qRT-PCR analysis of lncRNA expression in the right tibias of control (CON) and HLU mice (n=5). **(D)** The protein expression of Osx and Runx2 was determined by western blotting during osteoblast differentiation of MC3T3-E1 cells (n=3). **(E)** qRT-PCR analysis of Osx, Runx2 and OGRU RNA expression during osteoblast differentiation of MC3T3-E1 cells (n=3). \**P* < 0.05, \*\**P* <0.01.

### OGRU promotes the osteoblast activity and matrix mineralization and attenuates the osteogenic decline of MC3T3-E1 cells induced by clinorotation unloading

To assess the function of OGRU during osteoblast differentiation, MC3T3-E1 cells were transfected with pcDNA3.1(+)-OGRU, si-OGRU or the corresponding controls and were cultured in osteogenic medium. Notably, overexpression of OGRU in MC3T3-E1 cells significantly increased the mRNA levels of osteogenic marker genes, including Alp, Osx, Runx2 and Ocn (Figure 2A). Consistent with this finding, the protein expression levels of Osx, Runx2, and Ocn and Alp activity were increased in OGRU-overexpressing MC3T3-E1 cells (Figure 2B, C). Alp staining was significantly enhanced in the group treated with pcDNA3.1(+)-OGRU (Figure 2D). In addition, matrix mineralization, as visualized by alizarin red staining, was significantly promoted by pcDNA3.1(+)-OGRU (Figure 2E). In contrast, MC3T3-E1 cells transfected with si-OGRU exhibited impaired osteoblast activity and matrix mineralization. These data suggested that OGRU is a critical regulator of osteoblast activity and matrix mineralization in vitro under normal conditions. To explore whether forced expression of OGRU could rescue the osteogenic decline induced by unloading in vitro, MC3T3-E1 cells were transfected with pcDNA3.1(+)-OGRU for 12 h and were then cultured in clinorotation unloading condition for 48 h. These results showed that OGRU substantially attenuated the reduction in the mRNA levels of osteogenic marker genes, including Alp, Osx, Runx2 and Ocn, induced by clinorotation unloading (Figure 2F). Consistent with this result, the protein expression levels of Osx, Runx2, Ocn and Alp activity were also attenuated in OGRU-overexpressing MC3T3-E1 cells (Figure 2G, H).

**Figure 2.**
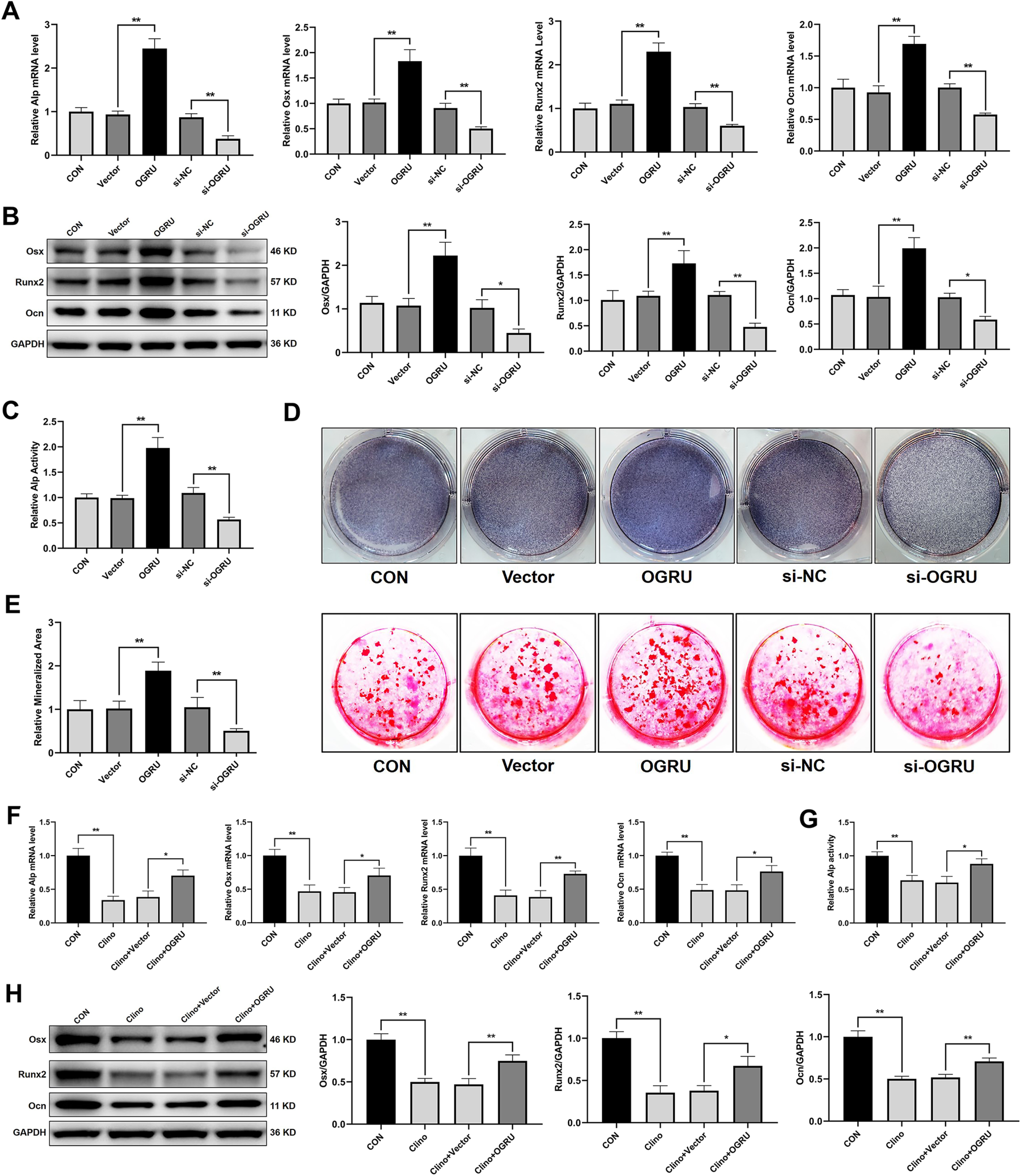
OGRU promotes the osteoblast activity and matrix mineralization and attenuates the osteogenic decline of MC3T3-E1 cells induced by clinorotation unloading. MC3T3-E1 cells were transfected with pcDNA3.1(+)-OGRU, si-OGRU or the corresponding controls and were cultured in osteogenic medium. **(A)** The mRNA levels of Alp, Osx, Runx2, and Ocn were measured by qRT-PCR 4 days after osteogenic treatment (n=3). **(B)** The protein levels of Osx, Runx2, and Ocn were measured by western blotting 4 days after osteogenic treatment (n=3). **(C)** The relative Alp activity 4 days after osteogenic treatment (n=3). **(D)** Representative images of Alp staining in MC3T3-E1 cells 7 days after osteogenic treatment (n=3). **(E)** Representative images of alizarin red staining in MC3T3-E1 cells 21 days after osteogenic treatment are shown, and relative areas of mineralization were quantified by Image J (n=3). MC3T3-E1 cells were transfected with pcDNA3.1(+) or pcDNA3.1(+)-OGRU and cultured in clinorotation unloading condition for 48 h and were then subjected to qRT-PCR analysis of Alp, Osx, Runx2 and Ocn mRNA levels **(F)**, measurement of the relative Alp activity **(G)** and western blot analysis of Osx, Runx2 and Ocn protein expression **(H)** (n=3). \**P* < 0.05, \*\**P* <0.01.

### Bone-targeted OGRU increases osteoblastic bone formation in HLU mice

As we found that OGRU played an important role in osteoblast activity and matrix mineralization under both normal and unloading conditions in vitro, we further explored whether therapeutic overexpression of OGRU could rescue unloading-induced bone loss in HLU mice. Thus, we preinjected pcDNA3.1(+)-OGRU carried by (DSS)_6_-liposome three times before HLU. After 21 days of HLU, mice were sacrificed, and the bilateral femurs and tibias were collected for bone analysis (Figure 3A). qRT-PCR analysis of RNA samples from mice tibias showed that bone-targeted OGRU effectively increased the expression of OGRU in bone (Figure 3B), which partially counteracted the decreased mRNA expression of osteogenic markers (Alp, Osx, Runx2 and Ocn) in HLU mice (Figure 3C-F). In addition, more Ocn-positive cells (osteoblasts) were found in the distal femurs from (DSS)_6_-liposome-OGRU-treated mice than in the femurs of (DSS)_6_-liposome-treated mice, as indicated by immunohistochemical staining (Figure 3D).

**Figure 3.**
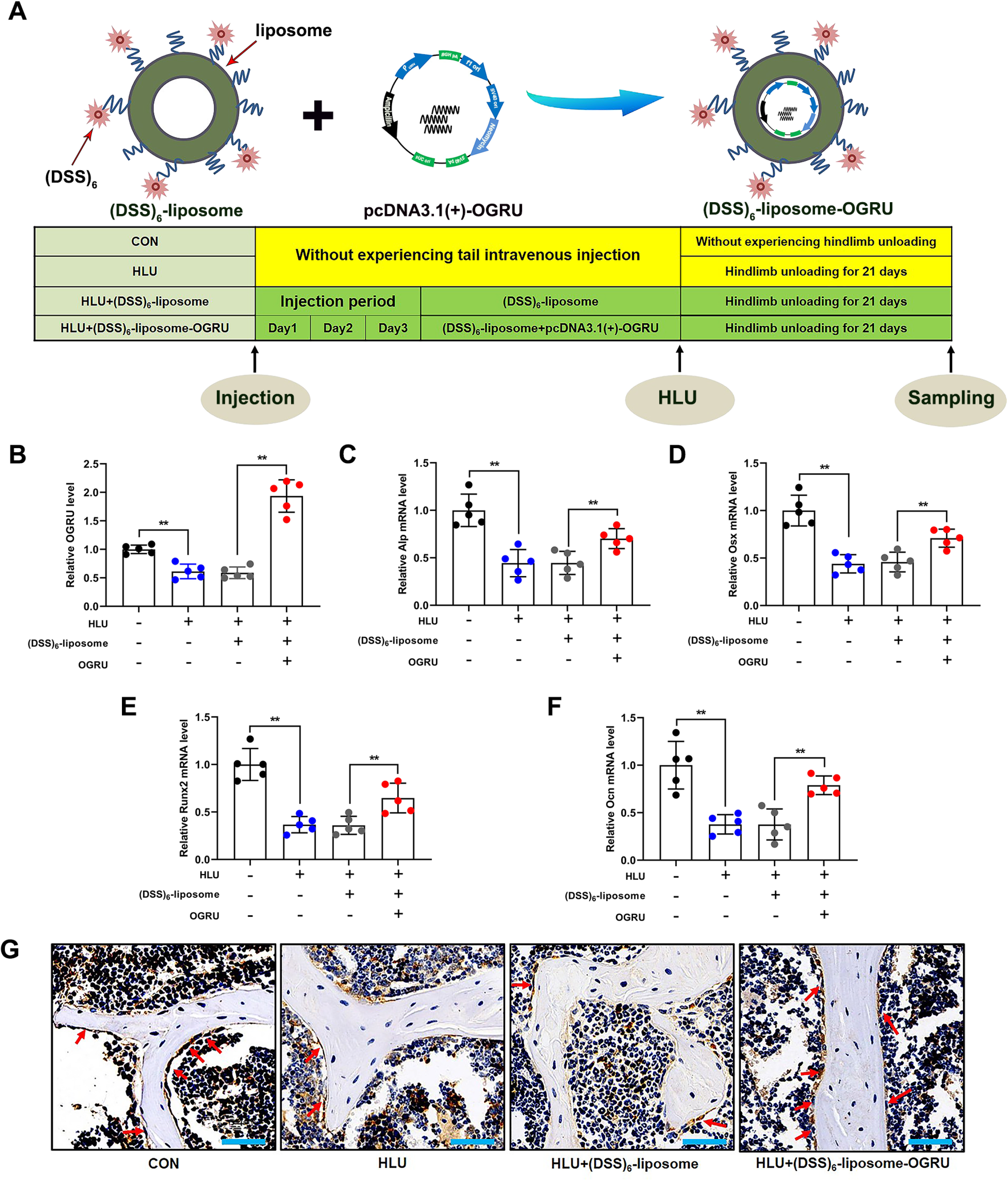
The effect of bone-targeted OGRU on osteogenic markers in HLU mice. **(A)** Schematic of the experimental design. **(B)** qRT-PCR analysis of OGRU expression in the right tibias of mice (n=5). **(C-F)** qRT-PCR detection of the Alp, Osx, Runx2 and Ocn mRNA levels in the right tibias of mice in the indicated groups (n=5). **(G)** Representative images of osteocalcin immunohistochemical staining in the right femurs of mice in the indicated groups (n=5). Scale bars: 50 μm. \*\**P* <0.01.

### Bone-targeted OGRU improves trabecular microarchitecture and increases bone mass in HLU mice

Next, we further explored the effects of bone-targeted OGRU on trabecular microarchitecture and bone mass. Micro-CT assays revealed that the significant decreases in bone mineral density (BMD), the ratio of bone volume to total volume (BV/TV), trabecular bone number (Tb.N) and trabecular thickness (Tb.Th) and the increases in the ratio of bone surface to bone volume (BS/BV) and trabecular bone pattern factor (TbPF) induced by HLU were efficiently attenuated in the (DSS)_6_-liposome-OGRU group (Figure 4A, B). H&E staining of the proximal side of the growth plate in the distal femurs further confirmed that (DSS)_6_-liposome-OGRU counteracted the low-bone-mass phenotype of HLU mice, as quantified by the ratio of bone area to total area (B.Ar/T.Ar) (Figure 4C). The data demonstrated that the low bone mass and deterioration of the trabecular microarchitecture in HLU mice was alleviated by bone-targeted OGRU.

**Figure 4.**
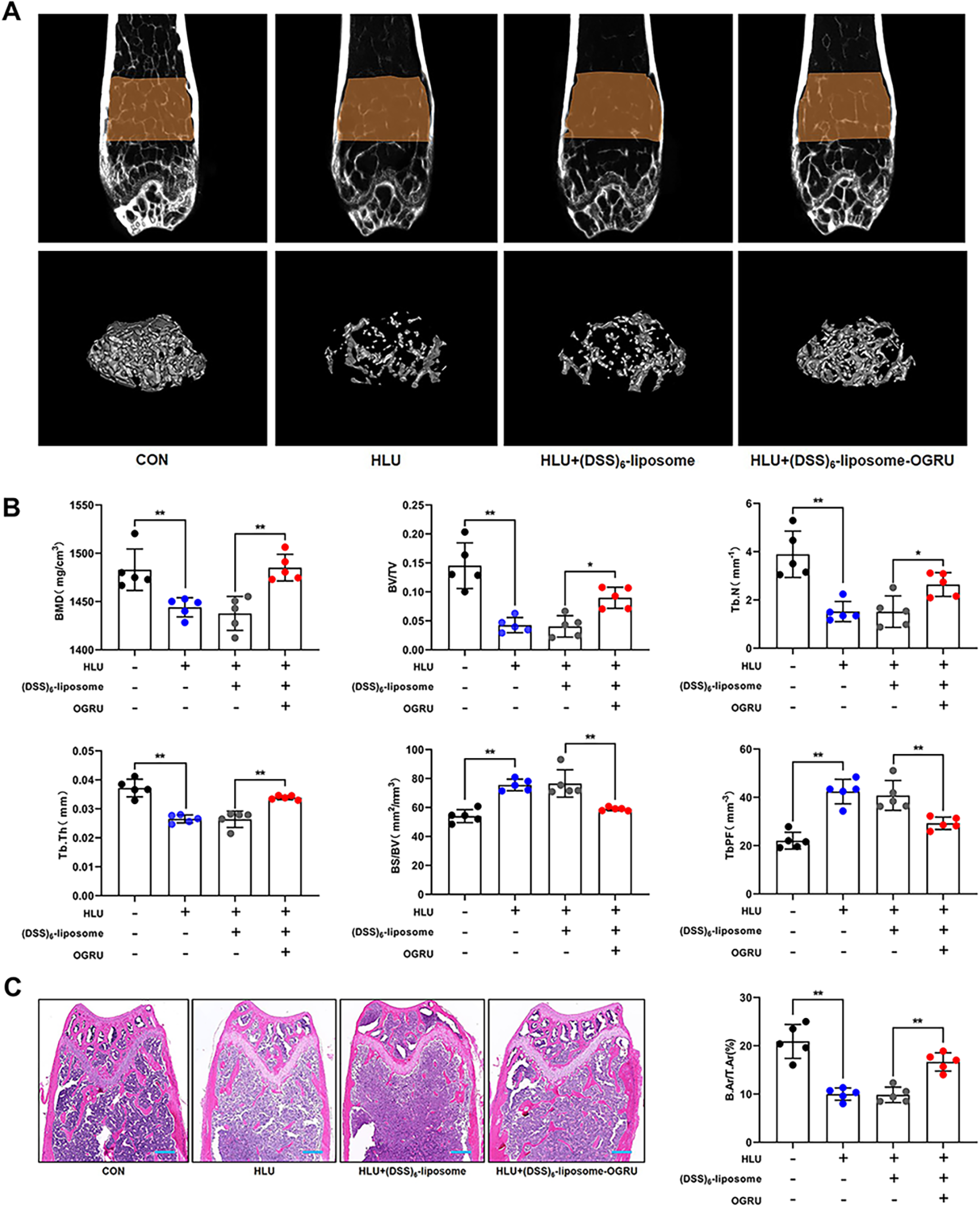
Bone-targeted OGRU improves trabecular microarchitecture and increases bone mass in HLU mice. **(A)** Representative images of micro-CT 2D sections and 3D reconstruction of distal femurs of mice in the indicated groups (n=5). **(B)** Micro-CT measurement of bone mineral density (BMD), the ratio of bone volume to total volume (BV/TV), trabecular bone number (Tb.N), trabecular thickness (Tb.Th), the ratio of bone surface to bone volume (BS/BV) and trabecular bone pattern factor (TbPF) (n=5). **(C)** H&E staining showing trabecular microarchitecture and quantitative analysis of the bone area/ total area on the proximal side of the growth plate in the distal femurs mice in the indicated groups (n=5). Scale bars: 300 μm. \**P* < 0.05, \*\**P* <0.01.

### Bone-targeted OGRU promotes new bone formation and improves biomechanical properties in HLU mice

To evaluate the role of bone-targeted OGRU in new bone formation in HLU mice, Goldner’s trichrome staining was performed and showed more newly formed bone in the distal femurs from (DSS)_6_-liposome-OGRU-treated mice than in the distal femurs from (DSS)_6_-liposome-treated mice (Figure 5A). Moreover, calcein double labeling analysis showed a higher osteogenic rate in the (DSS)_6_-liposome-OGRU group than in the (DSS)_6_-liposome group (Figure 5B).

**Figure 5.**
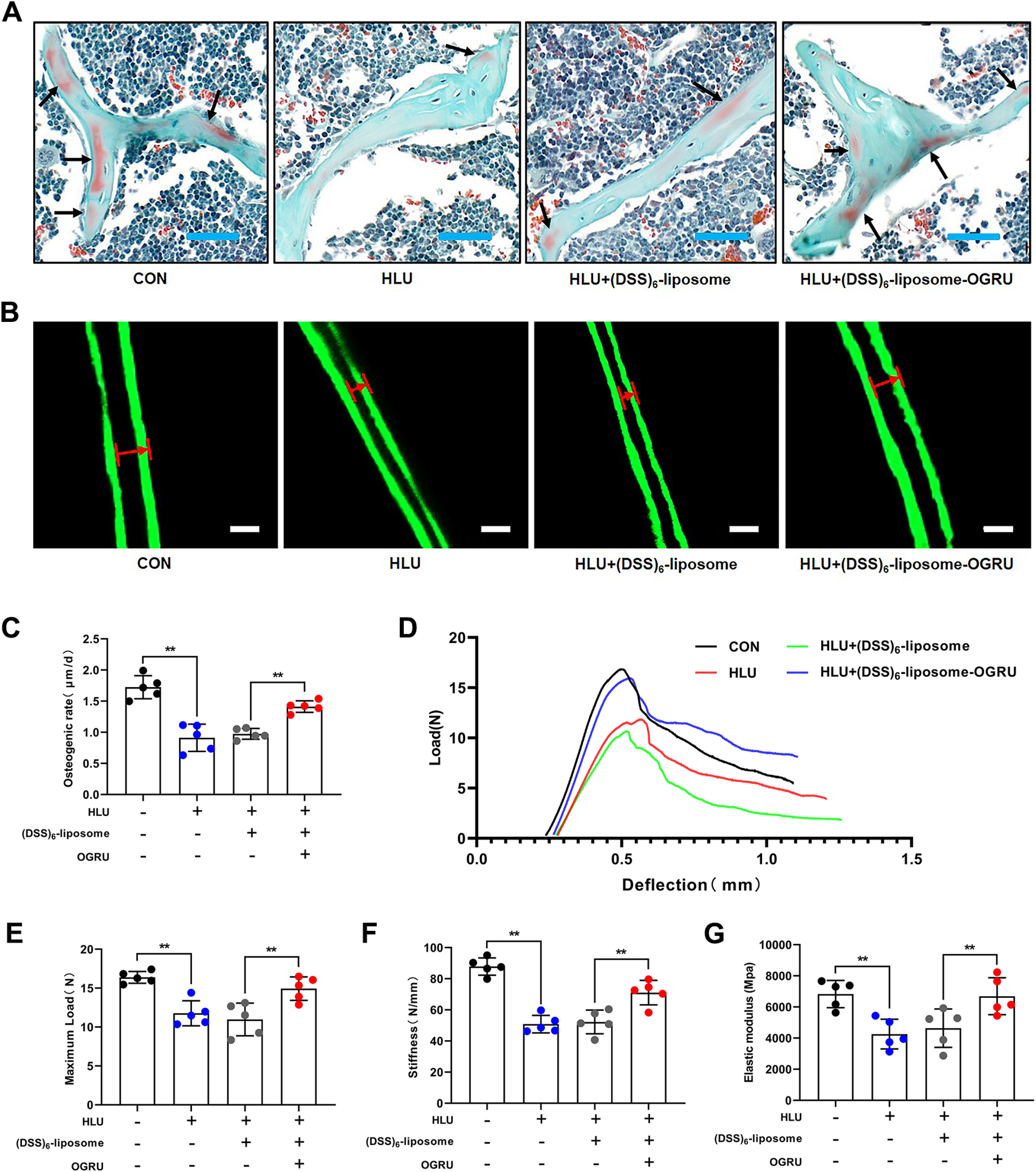
Bone-targeted OGRU promotes new bone formation and improves biomechanical properties in HLU mice. **(A)** Representative images of Goldner’s trichrome staining of the trabecular bone at distal femurs of mice in the indicated groups (n=5). Osteoid stains red (indicated by the black arrows), and mineralized bone stains green. Scale bars: 50 µm. **(B)** Representative images of calcein double labeling. The widths between the two fluorescence-labeled lines were measured with Image J (n=5). Scale bars: 10 μm. **(C)** Representative load-deflection curves for the respective groups (n=5). **(D)** The maximum load, **(E)** stiffness, and **(F)** elastic modulus were calculated (n=5). \*\**P* <0.01.

Biomechanical properties can indicate the resistance of bones to fracture, so we performed a three-point bending test on femurs to evaluate bone stiffness and strength. The structural parameters of the femurs, including the maximum load, stiffness, and elastic modulus, were calculated based on the load-deflection curves. The results showed that these three parameters were substantially decreased by HLU, but the impairment of these biomechanical properties was significantly ameliorated by (DSS)_6_-liposome-OGRU (Figure 5C-F).

### OGRU interacts with miR-320-3p

As the function of OGRU on osteogenesis during unloading was confirmed both in vitro and in vivo, we further explored the mechanisms involved in this process. The mechanism of lncRNA is closely related to its subcellular location, which was examined by RNA fluorescence in situ hybridization and cell fractionation followed by qRT-PCR, and the results showed that OGRU primarily localizes in the cytoplasm (Figure 6A, B). Thus, we speculated that OGRU may act as a ceRNA that binds to microRNAs. Bioinformatics analysis by RegRNA 2.0 (http://regrna2.mbc.nctu.edu.tw/) revealed that the OGRU sequence contains several putative binding sites for microRNAs, and we selected four miRNAs involved in osteoblast differentiation among these putative binding miRNAs for further study (Figure 6C, Table S5).^36–40^ The results demonstrated that the level of miR-320-3p was markedly increased after treatment with si-OGRU, but miR-320-3p had no effect on the expression of OGRU (Figure 6D, E). In addition, the miR-320-3p level was significantly increased during unloading in vitro and in vivo, and this effect was alleviated by OGRU overexpression (Figure 6F-H). Next, the binding between miR-320-3p and OGRU was further validated by dual luciferase reporter assays. We found that mimic-320 significantly reduced the luciferase activity of the reporter vector containing wild-type (WT) OGRU but not the reporter vector containing OGRU with mutated (MUT) miR-320 binding sites in 293T cells (Figure 6I, J). miRNAs can bind their targets in an Ago2-dependent manner. Therefore, to determine whether OGRU was regulated by miR-320-3p in such a manner, we conducted anti-Ago2 RNA-binding protein immunoprecipitation (RIP) in MC3T3-E1 cells and found that miR-320-3p and OGRU were enriched in the anti-Ago2 group compared to the anti-IgG group, as detected by qRT-PCR (Figure 6J). These data demonstrated that OGRU can act as a ceRNA for binding to miR-320-3p.

**Figure 6.**
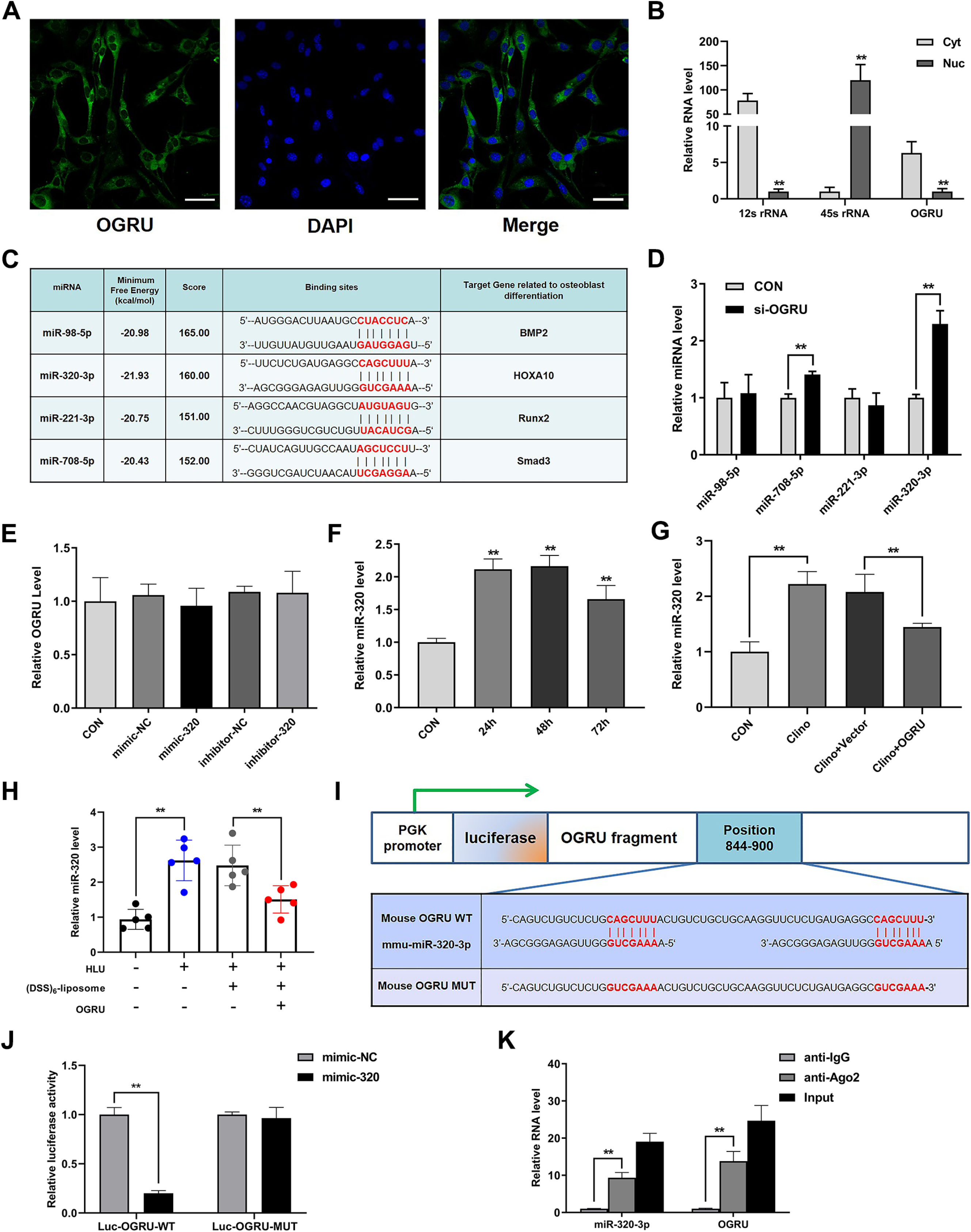
OGRU interacts with miR-320-3p. **(A)** RNA fluorescence in situ hybridization showed that OGRU primarily localizes in the cytoplasm (n=3). Scale bars: 50 µm. **(B)** qRT-PCR analysis of OGRU expression in the cytoplasmic (Cyt) and nuclear (Nuc) fractions. The control, 45S ribosomal RNA (rRNA), was primarily localized in the nucleus, and 12S rRNA was found primarily in the cytoplasmic fraction (n=3). **(C)** The RegRNA2.0 prediction for the binding sites for candidate miRNAs in the OGRU transcript. **(D)** qRT-PCR analysis of miRNA levels after treatment with si-OGRU in MC3T3-E1 cells (n=3). **(E)** qRT-PCR analysis of OGRU expression levels after treatment with mimic-320, inhibitor-320 or the corresponding controls (n=3). **(F)** qRT-PCR measurement of the relative miR-320-3p levels in MC3T3-E1 cells under clinorotation unloading condition for 24, 48 and 72 h (n=3). **(G)** qRT-PCR measurement of miR-320-3p levels after MC3T3-E1 cells were transfected with pcDNA3.1(+)-OGRU and subjected to clinorotation unloading for 48 h (n=3). **(H)** qRT-PCR detection of the miR-320-3p levels in the right tibias from mice in the CON, HLU, HLU+(DSS)_6_-liposome and HLU+(DSS)_6_-liposome-OGRU groups (n=5). **(I, J)** The relative luciferase activity of luciferase reporters containing OGRU wild-type (WT) or mutated (MUT) miR-320 binding sites in 293T cells cotransfected with mimic-320 or its negative control was measured. The data are presented as the relative ratio of firefly luciferase activity to renilla luciferase activity (n=3). **(K)** Anti-AGO2 RIP was performed using input from cell lysate, normal mouse IgG or anti-Ago2, followed by qRT-PCR to measure the OGRU and miR-320 levels (n=3). \**P* < 0.05, \*\**P* <0.01.

### miR-320-3p inhibits the osteoblast activity and matrix mineralization of MC3T3-E1 cells by targeting Hoxa10

A previous study confirmed that hsa-miR-320a-3p inhibits the osteoblast activity and matrix mineralization of human bone marrow-derived mesenchymal stem cells (hMSCs) by targeting Hoxa10, a critical regulator of osteogenesis.^37, 41, 42^ Considering that mmu-miR-320-3p is the murine homolog of hsa-miR-320a-3p and was predicted to target Hoxa10 by TargetScan, miRanda, and miRDB (Table S6), we explored whether miR-320-3p could regulate the osteoblast activity and matrix mineralization of MC3T3-E1 cells in such a manner. During the osteoblast differentiation phase, miR-320-3p levels decreased gradually, whereas Hoxa10 protein levels increased (Figure 7A, B). Moreover, miR-320-3p significantly influenced Hoxa10 protein expression but had no effect on Hoxa10 mRNA levels (Figure 7C, D). Next, we constructed luciferase reporters containing either the WT Hoxa10 3′UTR or the Hoxa10 3′UTR with mutated (MUT) miR-320 binding sites in 293T cells and found that mimic-320 substantially inhibited the luciferase reporter activity of the WT Hoxa10 3′UTR but not the MUT Hoxa10 3′UTR (Figure 7E). Taken together, our results confirmed that Hoxa10 is the target of miR-320-3p.

**Figure 7.**
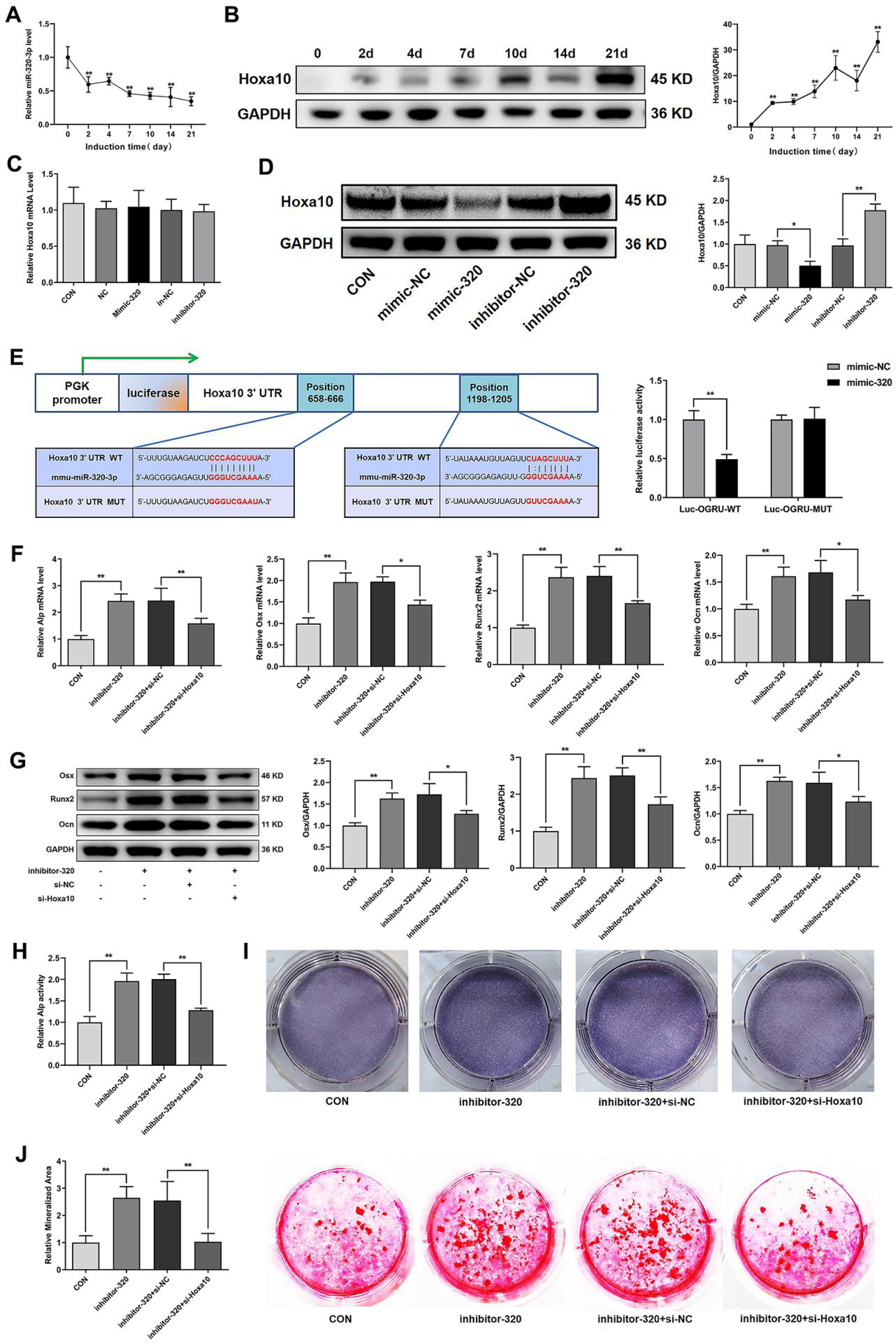
miR-320-3p inhibits the osteoblast activity and matrix mineralization of MC3T3-E1 cells by targeting Hoxa10. **(A)** qRT-PCR analysis of miR-320-3p levels during osteoblast differentiation of MC3T3-E1 cells (n=3). **(B)** Western blot analysis of Hoxa10 protein expression during osteoblast differentiation of MC3T3-E1 cells (n=3). **(C)** qRT-PCR analysis of Hoxa10 mRNA expression levels and **(D)** western blot analysis of Hoxa10 protein expression levels after treatment with mimic-320, inhibitor-320 or the corresponding controls (n=3). **(E)** The relative luciferase activity of luciferase reporters containing the wild-type (WT) Hoxa10 3’UTR or the Hoxa10 3’UTR with a mutated (MUT) miR-320 binding site in 293T cells cotransfected with mimic-320 or its negative control was measured (n=3). MC3T3-E1 cells were cotransfected with inhibitor-320 and si-Hoxa10 or its negative controls. qRT-PCR analysis of Alp, Osx, Runx2 and Ocn mRNA levels **(F)**, western blot analysis of Osx, Runx2 and Ocn protein levels **(G)**, relative Alp activity **(H)**, representative images of Alp staining in MC3T3-E1 cells **(I)**, and representative images of alizarin red staining in MC3T3-E1 cells are presented, and the relative areas of mineralization were quantified by Image J **(J)** (n=3). \**P* < 0.05, \*\**P* <0.01.

To determine whether Hoxa10 is responsible for the regulatory effect of miR-320-3p on osteoblast activity and matrix mineralization, MC3T3-E1 cells were cotransfected with inhibitor-320 and si-Hoxa10 or its negative control and were cultured in osteogenic medium. The results showed that si-Hoxa10 partially reduced the promotion of osteoblast activity and matrix mineralization induced by inhibitor-320, as determined by the mRNA or protein expression of osteogenic marker genes, Alp activity, Alp staining and matrix mineralization (Figure 7F-J).

### Hoxa10 is responsible for OGRU-mediated osteoblast differentiation and bone formation during unloading

Since OGRU interacted with miR-320-3p and Hoxa10 was the target of miR-320-3p, we examined whether OGRU could positively regulate the expression of Hoxa10. Our results demonstrated that OGRU promoted the protein expression of Hoxa10 and partially reversed the reduction in Hoxa10 protein expression induced by clinorotation unloading in MC3T3-E1 cells but did not influence the mRNA expression of Hoxa10 (Figure 8A-D). Immunohistochemical staining of Hoxa10 also indicated that OGRU partially reversed the reduction in Hoxa10 protein expression induced by HLU (Figure 8E). In addition, the effect of the enhanced osteoblast differentiation induced by OGRU during unloading was reversed by si-Hoxa10, as evidenced by the mRNA levels of Alp, Osx, Runx2 and Ocn; the protein expression of Osx, Runx2, and Ocn; and Alp activity (Figure 8F-H).

**Figure 8.**
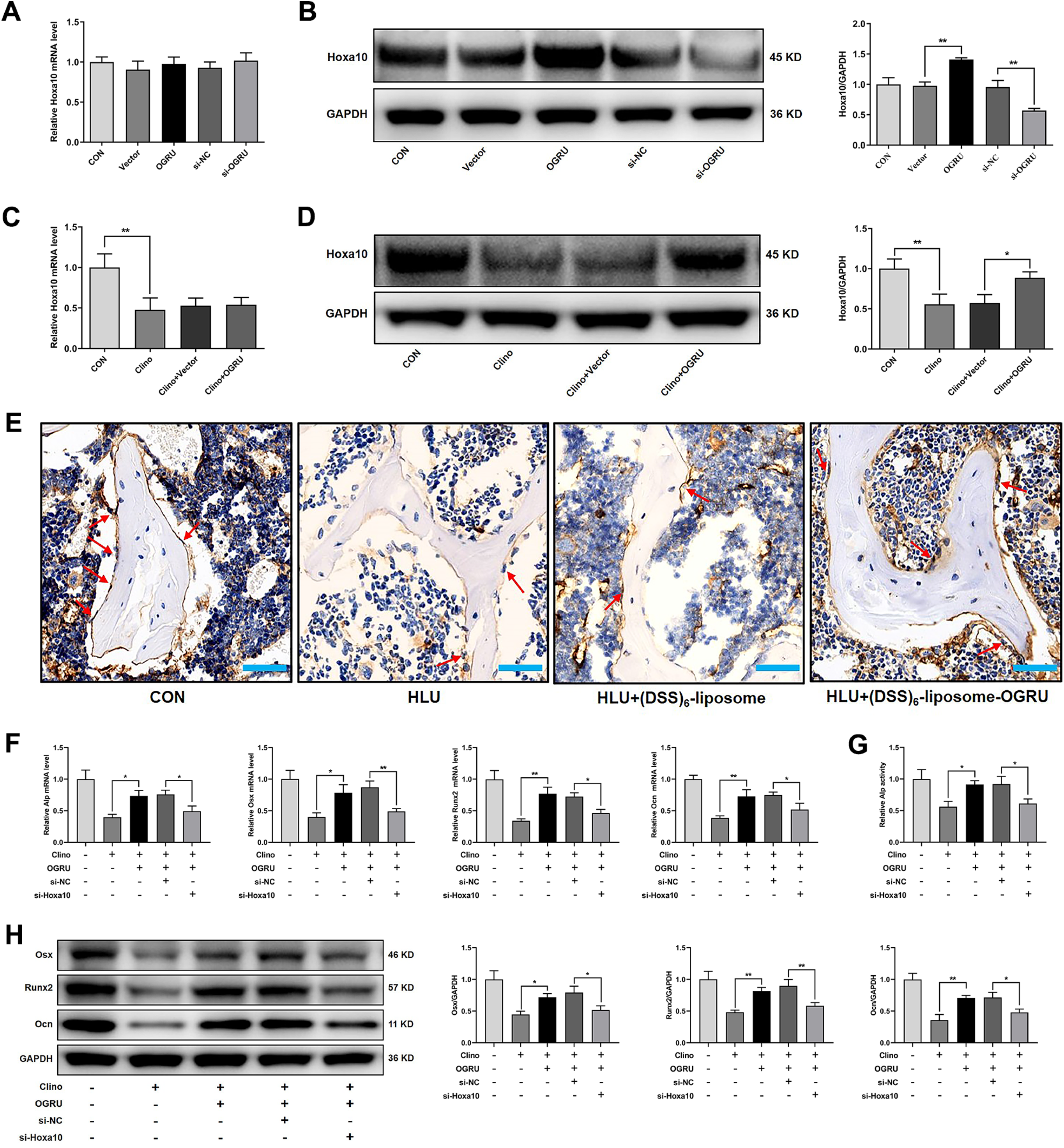
Hoxa10 is responsible for OGRU-mediated osteoblast differentiation and bone formation during unloading. **(A)** qRT-PCR analysis of Hoxa10 mRNA levels and **(B)** western blot analysis of Hoxa10 protein levels after treatment with pcDNA3.1(+)-OGRU, si-OGRU or the corresponding controls in MC3T3-E1 cells (n=3). **(C)** Hoxa10 mRNA levels were measured by qRT-PCR, and **(D)** Hoxa10 protein expression was measured by western blotting after MC3T3-E1 cells were transfected with pcDNA3.1(+) or pcDNA3.1(+)-OGRU and subjected to clinorotation unloading for 48 h (n=3). **(E)** Representative images of Hoxa10 immunohistochemical staining in the right femurs of mice from the indicated groups (n=5). Scale bars: 50 µm. **(F)** qRT-PCR analysis of Alp, Osx, Runx2 and Ocn mRNA levels, **(G)** western blot analysis of Osx, Runx2, Ocn protein expression and **(H)** the relative Alp activity after MC3T3-E1 cells were cotransfected with pcDNA3.1(+)-OGRU and si-Hoxa10 or its negative controls and subjected to clinorotation unloading for 48 h (n=3). \**P* < 0.05, \*\**P* <0.01.

**Figure 9.**
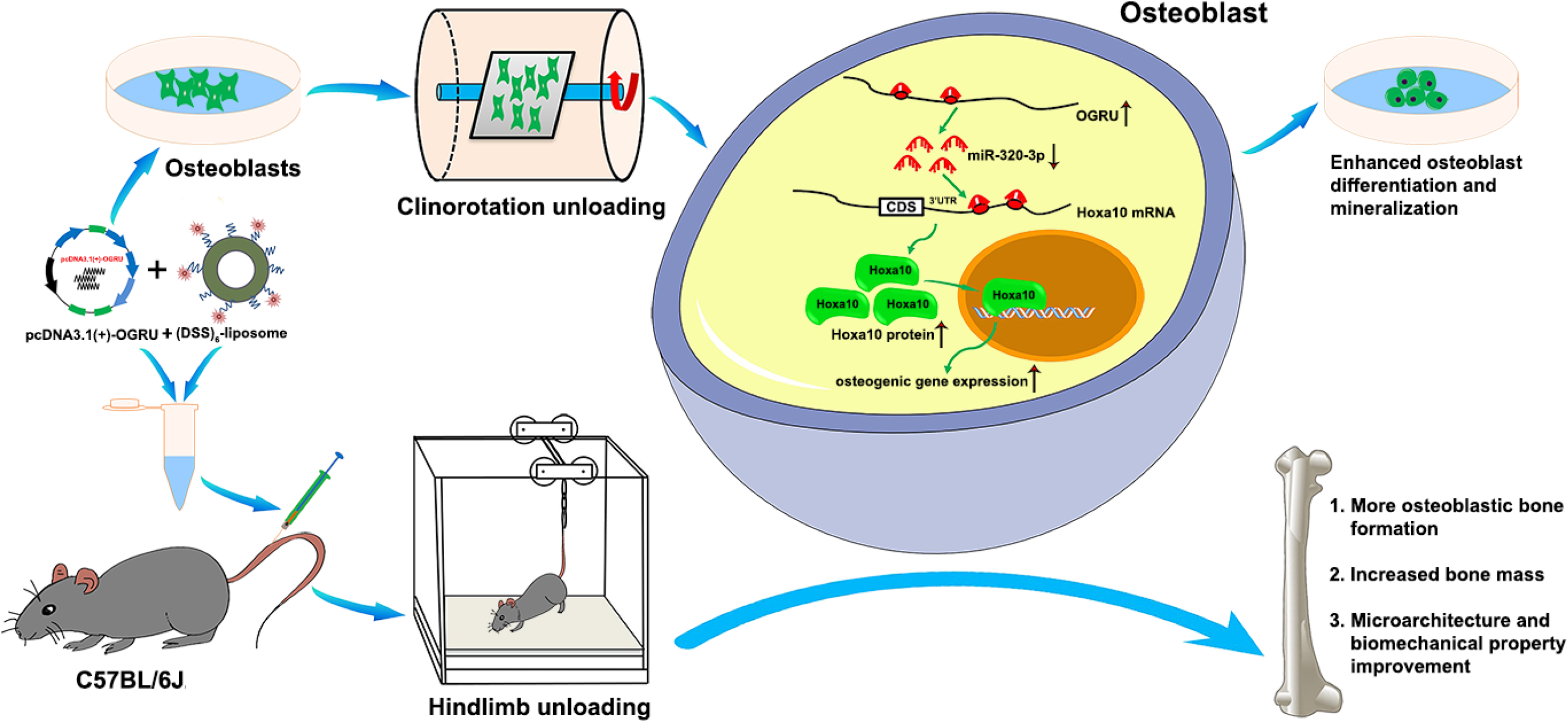
A Schematic illustrating the molecular mechanisms of OGRU-regulated unloading-induced bone loss: OGRU can significantly promote Hoxa10 protein expression by functioning as a miR-320-3p sponge, which results in enhanced osteoblast differentiation and mineralization in vitro and more osteoblastic bone formation, increased bone mass and trabecular microarchitecture and biomechanical properties improvement via (DSS)_6_-liposome delivery system in vivo during mechanical unloading condition.

## Discussion

Skeletal unloading by long-duration space flight and extended bed rest can induce disuse osteopenia, which is regulated by various cytokines, hormones and miRNAs.^14, 43, 44^ However, scant information addresses the role of lncRNAs in unloading-induced bone loss. In the present study, we identified a novel unloading-sensitive lncRNA, OGRU, and confirmed its direct impact on osteoblast differentiation during unloading in vitro. Moreover, bone-targeted OGRU counteracted bone loss in HLU mice. The mechanistic studies showed that OGRU sponges miR-320-3p to promote osteoblast differentiation and bone formation by upregulating the protein expression of Hoxa10. Thus, our research was the first to elucidate the role of the OGRU/miR-320-3p/Hoxa10 pathway in the pathophysiological process of unloading-induced bone loss.

Multiple lncRNAs have been identified as important regulators of osteoblast differentiation and bone formation via a variety of mechanisms dependent on their subcellular location.^45^ For instance, the lncRNA Bmncr can alleviate bone loss and bone marrow fat accumulation during aging by facilitating the assembly of the TAZ and RUNX2/PPARG transcriptional complex.^29^ Moreover, the lncRNA MEG3 can activate BMP4 transcription by dissociating SOX2 from the BMP4 promoter and thereby promote osteoblast differentiation of MSCs.^28^ An additional study reported that the lncRNA PGC1β-OT1 can stimulate osteoblast differentiation of progenitor cells in vitro and in vivo by antagonizing the function of miR-148a-3p.^46^ In particular, the lncRNA H19 (H19) is a key regulator of osteoblast function and is involved in the development of disuse osteoporosis, suggesting that lncRNAs may play a key role in unloading-induced bone loss.^31, 32, 47, 48^ Our previous work found many lncRNAs that are significantly differentially expressed in MC3T3-E1 cells under clinorotation unloading condition for 48 h. Further exploration identified a novel unloading-sensitive lncRNA, OGRU, which promotes the osteoblast activity and matrix mineralization and attenuates the osteogenic impairment of MC3T3-E1 cells induced by clinorotation unloading. To further determine whether OGRU can rescue unloading-induced bone loss in vivo, pcDNA3.1(+)-OGRU was delivered via (DSS)_6_-liposomes to bone formation surfaces in HLU mice. In this regard, OGRU can promote bone formation and can be used as a therapeutic target against unloading-induced bone loss. However, it has been reported that unloading-induced bone loss is caused by both decreased bone formation and increased bone resorption. In this study, OGRU was delivered only to bone formation surfaces in HLU mice; thus, its effects on bone resorption have not been elucidated.

To date, many RNA transcripts, such as lncRNAs and circular RNAs, have been thoroughly characterized as ceRNAs for microRNA binding.^49, 50^ Here, we found that OGRU contains binding sites for and can interact with miR-320-3p in the Ago2-RISC complex. Previously, it has been reported that miR-320 is a critical regulator of tumorigenesis, inflammation and substance metabolism.^51–54^ In addition, hsa-miR-320a-3p inhibits osteoblast differentiation of hMSCs.^37^ However, scant information addresses the role of mmu-miR-320-3p in osteoblast differentiation and bone formation, especially during unloading. In this study, in vitro and in vivo experiments strongly suggested that miR-320-3p is unloading-sensitive and that its levels can be decreased by OGRU during unloading. Furthermore, we demonstrated that miR-320-3p inhibits the osteoblast activity and matrix mineralization of MC3T3-E1 cells by targeting Hoxa10.

Hoxa10, a member of the homeobox gene family, can participate in the development and growth of multiple systems, as indicated by its critical roles in male and female fertility, liver tumorigenesis, gastric cancer invasion and leukemogenesis.^55–58^ Moreover, multiple lines of evidence have shown that Hoxa10 is necessary for the global patterning of the mammalian skeleton.^59–61^ Furthermore, Hoxa10 can promote osteoblast differentiation through the activation of Runx2 and directly regulate osteoblast differentiation genes, including the alkaline phosphatase, osteocalcin, and bone sialoprotein genes, and this process can be regulated by the Pbx1 complex.^41, 42, 62, 63^ In ovariectomized mice, Hoxa10 protein levels were significantly reduced by miR-705, but Hoxa10 mRNA levels were not significantly changed.^64^ Here, we reported that both the protein and mRNA levels of Hoxa10 were decreased by unloading in vitro and in vivo, but only the decrease in Hoxa10 protein levels was ameliorated by the OGRU/miR-320-3p axis. Accordingly, we speculated that Hoxa10 can also be regulated by other mechanisms at the transcriptional and epigenetic levels during unloading.

In summary, our findings uncover a novel unloading-sensitive lncRNA, OGRU, which is a critical regulator of osteoblast activity and matrix mineralization and attenuates osteogenic decline by functioning as a miR-320-3p sponge to stimulate Hoxa10 protein expression. Correspondingly, bone-targeted OGRU can effectively counteract unloading-induced bone loss, indicating that OGRU may be a promising therapeutic target for unloading-induced bone loss and disuse osteoporosis.

## Materials and Methods

### Cell culture and in vitro differentiation

The murine preosteoblast cell line MC3T3-E1 clone 14 was purchased from the Cell Bank of the Chinese Academy of Sciences (Shanghai, China) and cultured in alpha modified Eagle’s medium (α-MEM, Gibco, USA) supplemented with 10% FBS (HyClone, USA) and 1% penicillin and streptomycin (Invitrogen, USA) in an atmosphere of 5% CO_2_ and 95% humidity at 37 °C. The culture medium was changed every 2 days. Cells beyond passage 15 were not used for experiments. For osteoblast differentiation, MC3T3-E1 cells at an appropriate confluency were induced by the addition of 100 nM dexamethasone, 10 mM β-glycerophosphate, and 50 μg/ml ascorbic acid to the culture medium.

### 2D clinorotation

A 2D clinostat (developed by the China Astronaut Research and Training Center, Beijing, China) was selected to simulate the unloading environment in vitro, and related experiments were performed according to the procedure described previously.^13, 65^ Briefly, MC3T3-E1 cells were seeded in 25-cm^2^ culture flasks at a density of 5×10^5^ cells per flask. After cell adherence, the culture flasks were filled with osteogenic medium, ensuring that air bubbles were removed. Then, the culture flasks were fixed to the 2D clinostat and rotated around the horizontal axis at 24 rpm for 24, 48, and 72 hours. The CON groups were placed in a similar incubator without clinorotation.

### HLU mice and experimental design

Male C57BL/6J mice purchased from the Animal Center of Air Force Medical University (Xi’an, China) were housed under standard conditions (22 °C, 50%-55% humidity, and a 12 h light/12 h dark cycle). HLU was performed to obtain a mouse model with unloading-induced bone loss, as previously reported.^66^ Briefly, 6-month-old mice were suspended at a 30° angle with the floor by the tail, which allowed the mice to move and access food and water freely. As illustrated in Figure 3A, twenty 6-month-old male C57BL/6J mice were randomly divided into four groups, as follows: (1) CON, (2) HLU, (3) HLU+(DSS)_6_-liposome, and (4) HLU+(DSS)_6_-liposome-OGRU (n = 5 per group). Before HLU, either pcDNA3.1(+)-OGRU (2 mg/kg body weight) carried by (DSS)_6_-liposomes or (DSS)_6_-liposomes alone was administered by three consecutive intravenous injections. After 21 days of HLU, the bilateral femurs and tibias of the mice were collected for bone analysis. All experimental protocols were approved by the Animal Care Committee of Air Force Medical University.

### RNA extraction and quantitative real-time polymerase chain reaction (qRT-PCR) analysis

Total RNA was isolated using RNAiso Plus according to the manufacturer’s protocol (TaKaRa, China). Total RNA was reverse transcribed to complementary DNA (cDNA) using a Prime Script™ RT Master Mix Kit (TaKaRa, China) and the following procedure: 37 °C for 15 minutes, 85 °C for 5 seconds, and holding at 4 °C. For miRNA expression analysis, miRNA was reverse transcribed using a Mir-X miRNA First-Strand Synthesis Kit (Clontech, USA) and the following procedure: incubation at 37 °C for 1 hour, termination at 85 °C for 5 minutes, and holding at 4 °C. qRT-PCR reactions were performed using a SYBR® Premix Ex Taq™ II Kit (TaKaRa, China) and a CFX96 real-time PCR detection system (BIO-RAD, USA). The expression of lncRNAs and mRNAs was analyzed using the following cycling program: denaturation for 2 minutes at 95 °C, 40 cycles of annealing for 10 seconds at 95 °C and elongation for 40 seconds at 60 °C. GAPDH was used as the reference gene. The expression of miRNA was analyzed using the following cycling program: 30 seconds at 95 °C, 40 cycles of 5 seconds at 95 °C and 30 seconds at 60 °C. U6 was used as the reference gene. The primer sequences used are shown in Table S1.

### Protein isolation and western blotting analysis

Total protein was extracted from cells using M-PER Mammalian Protein Extraction Reagent (Thermo Scientific, USA) supplemented with protease inhibitor cocktail (Roche Applied Science, Germany) and was quantitated by a Pierce™ BCA Protein Assay Kit (Thermo Scientific, USA) following the manufacturer’s protocol. After protein samples were heated at 70 °C for 10 minutes, appropriate amounts of samples containing protein along with LDS sample buffer, reducing agent (Thermo Scientific, USA) and double-distilled H_2_O were subjected to electrophoresis on NuPAGE™ Bis-Tris Protein Gels (Invitrogen, USA). After electrophoresis, the separated proteins were transferred to polyvinylidene difluoride (PVDF) membranes with a constant current of 250 mA for 2 hours at 4°C. Then, PVDF membranes were blocked with 5% nonfat milk in TBST for 2 hours at room temperature and incubated with primary antibodies specific for Osx (1:1000, Abcam, UK), Runx2 (1:1000, Cell Signaling Technology, USA), Ocn (1:2000, Abcam, UK), GAPDH (1:5000, Proteintech, USA), and Hoxa10 (1:1000, Santa Cruz, USA) in primary antibody dilution buffer (Beyotime, China) overnight at 4 °C. Next, PVDF membranes were incubated with peroxidase-conjugated goat anti-rabbit IgG or goat anti-mouse IgG (1:5000, ZSGB-BIO, China) for 1 hours at room temperature and were visualized by chemiluminescence reagent (Millipore, USA). GAPDH served as the reference gene. The relative quantity of protein expression was analyzed with the Image J software.

### 5’ and 3’ Rapid Amplification of cDNA Ends (RACE)

The transcription initiation and termination sites of OGRU were detected by 5’ and 3’ RACE using a SMARTer® RACE 5’/3’ Kit (Clontech, USA) according to the manufacturer’s instructions. Briefly, RNA was extracted from MC3T3-E1 cells, and 3’-and 5’-RACE-Ready cDNA was synthesized using SMARTScribe Reverse Transcriptase, as follows: 42 °C for 90 minutes and 70 °C for 10 minutes. Amplification was performed as follows: 5 cycles at 94 °C for 30 seconds and 72 °C for 3 minutes; 5 cycles at 94 °C for 30 seconds, 70 °C for 30 seconds, and 72 °C for 3 minutes; and 20 cycles at 94 °C for 30 seconds, 68 °C for 30 seconds, and 72 °C for 3 minutes. The obtained gel products were extracted and cloned into the pEASY cloning vector for sequencing. The gene-specific primers used are listed in Table S2.

### Northern blot analysis

Total RNA extracted from cells combined with 5× formaldehyde gel electrophoresis buffer, 37% formaldehyde and formamide were heated at 65 °C for 15 minutes and cooled on ice for 5 minutes. RNA loading buffer was then added. Samples were subjected to formaldehyde gel electrophoresis at 50 V for 2 hours and transferred to a nylon membrane. After prehybridization for 3 h at 42 °C, the membrane was hybridized with a DIG-labeled probe complementary to OGRU at 42 °C for 16 hours. The membrane was visualized using a Tanon 4600 (Tanon, China). The DIG-labeled probe sequence complementary to OGRU was as follows: TTTGGTTGACTTCCCTGATACTTCAGAAAGATAAGAAAATGAACTCTACTCTCTTGCTTCTGGATCTTTTGTTCCCCTCTGTCTCCCCATTCCTTTCCTCCAACTCTCCACATGTTAATGGCTGGCCTCTCCTTATCTACTCTTTCTCTCTGCCTTTCTCGACTCTAGGACCCTCTTAA CTC.

### Cell transfection

Cells were transfected with the miR-320-3p mimic (40 nM) and miR-320-3p inhibitor (80 nM) or their corresponding negative controls (GenePharma, China) using a Lipofectamine 2000 kit (Invitrogen, USA) following the manufacturer’s instructions. siRNAs (80 nM) specific for lnc-OGRU or Hoxa10 (GenePharma, China) and pcDNA3.1(+)-OGRU (200 ng/μl) (GeneCreate, China) were transfected into cells using a Lipofectamine 3000 kit (Invitrogen, USA). The sequences of the siRNAs are shown in Table S3.

### Alkaline phosphatase activity assay

Total protein extracted from cells was utilized to measure Alp activity using an alkaline phosphatase assay kit (Nanjing Jiancheng Technological, China) following the manufacturer’s protocol. The protein concentration was measured by the Pierce™ BCA Protein Assay Kit (Thermo Scientific, USA). Phenol (0.02 mg/ml) was used as the standard solution, and double-distilled H_2_O was used as the blank control solution. Alp activity was defined as the amount of phenol produced after 1 g of protein was reacted with the substrate for 15 minutes at 37 °C.

### Alkaline phosphatase staining

After MC3T3-E1 cells were cultured in osteogenic medium for 7 days, alkaline phosphatase staining was performed. Briefly, cells in 6-well plates were fixed with 4% paraformaldehyde for 15 minutes and stained with a BCIP/NBT Alp Color Development kit (Beyotime, China) for 30 minutes at room temperature according to the manufacturer’s instructions. The whole procedure was performed in the dark. Images were acquired by a digital camera.

### Alizarin red staining

MC3T3-E1 cells cultured in osteogenic medium for 21 days were washed with DPBS three times and fixed with ice-cold 70% ethanol for 40 minutes at 4 °C. After three washes with double-distilled H_2_O, cells were stained in 1% alizarin red S (Sigma-Aldrich, USA) for 15 minutes at room temperature. Next, cells were rinsed five times with double-distilled H_2_O, and the stained cells were imaged by a digital camera. The relative areas of mineralization were quantified by Image J software.

### Micro-CT analysis

Dissected right femurs were fixed with 4% paraformaldehyde for 2 days and scanned with a micro-CT imaging system (Siemens, Germany) with a spatial resolution of 10.44 μm (80 kV, 500 mA, 800 ms integration time) as previously reported.^44^ The region of interest (ROI) selected was a 2.5 × 2.5 × 3 mm^3^ cube approximately 1.5 mm from the growth plate in the distal femurs. A 3D reconstruction of the ROI was used to determine the BMD, BV/TV, BS/BV, Tb.Th, Tb.N and TbPF values with COBRA software.

### Histological analysis

After micro-CT analysis, the right femurs were decalcified in 10% EDTA for 3 weeks and embedded in paraffin before sectioning (5 μm, longitudinally oriented). Bone sections were subjected to H&E and Goldner’s trichrome staining according to the standard protocol. For immunohistochemical staining, the primary antibodies used were specific for Ocn (1:100, Abcam, UK) or Hoxa10 (1:100, Santa Cruz, USA). Sections were examined with a microscope (Eclipse E100, Nikon).

### Double calcein labeling assay

Mice received intraperitoneal injection of calcein (8 mg/kg body weight, Sigma-Aldrich, USA) 10 days and 3 days before sacrifice. After the collected right tibias were fixed with 4% paraformaldehyde for 2 days and embedded in polymethyl acrylate, the samples were serially cut into 50 μm thick sections with a hard tissue slicing machine (SP1600, Leica, Germany) in the dark, and double calcein labeling was imaged by confocal microscopy (LSM800, ZEISS). The distance between two fluorescence-labeled lines as measured with Image J software was used to evaluate the osteogenic rate.

### Biomechanical testing

The dissected right femurs were wrapped with gauze dipped in saline and stored in −80 °C. A three-point bending test was performed using an electromechanical material testing machine (Bose, USA). The femurs were placed on a supporter with two points with an 8 mm span distance, and a load was applied vertically to the midshaft of the femurs at a constant displacement rate of 0.02 mm/s until fracture. Then, the lengths of the external major axis and minor axis and the thickness of the cortical bone at the fracture site were measured using a Vernier caliper. The values of the structural properties were calculated according to the load-deflection curve and included the maximum load, stiffness, and elastic modulus.^67^

### RNA Fluorescence In Situ Hybridization

Cells were fixed with 4% paraformaldehyde for 20 minutes at room temperature. Then, hybridization solution containing a FAM-labeled OGRU probe (5’-TGAAGTATCAGGGAAGTCA ACCAA-3’) was added, and cells were hybridized overnight at 37 °C. DAPI was used to stain nuclei. Images were acquired under a fluorescence microscope (Eclipse E100, Nikon).

### Nuclear-cytoplasmic fractionation

Cytoplasmic and nuclear RNA were isolated from MC3T3-E1 cells with a PARIS™ Kit (Invitrogen, USA) following the manufacturer’s instructions. RNA samples were reverse transcribed to cDNA, and qRT-PCR was performed to detect OGRU expression in the nucleus and cytoplasm, as described above. 45S rRNA (primarily in the nucleus) and 12S rRNA (primarily in the cytoplasm) were used as controls.

### Plasmid constructs and luciferase activity assays

The luciferase constructs containing OGRU-WT and Hoxa10 3’UTR WT or OGRU-MUT and Hoxa10 3’UTR MUT were generated by inserting a PCR fragment containing the predicted or mutated binding sites of miR-320-3p into the pmirGLO vector between the SacI and XhoI sites. The luciferase construct and the miR-320-3p mimic or its negative control were cotransfected into 293T cells. Luciferase activity was measured 24 hours post transfection using the Dual Luciferase Reporter Assay System (Promega, USA). Firefly luciferase activity was normalized to renilla luciferase activity.

### RNA-binding protein immunoprecipitation (RIP)

RIP experiments were performed using a Magna RIP™ RNA-Binding Protein Immunoprecipitation Kit (Millipore, USA) according to the manufacturer’s instructions. The antibodies used were anti-Ago2 (Abcam, UK) and anti-IgG (Millipore, USA). qRT-PCR was used to detect OGRU and miR-320-3p expression among the precipitated RNAs.

### Statistical analysis

All data are expressed as the means ± SD and were analyzed using SPSS Statistics 22.0. Two-group comparisons were performed using Student’s t-test, and multiple group comparisons were analyzed by one-way ANOVA followed by the LSD post hoc test. *P* < 0.05 was considered statistically significant.

## Acknowledgments

This work was supported by grants from the National Natural Science Foundation of China (grant nos. 31570939, 81701856), the Key Pre-research Project of Manned Spaceflight (grant no. 020106), the Key Research and Development Program of Shaanxi (Program No. 2018SF-039) and Young Talent fund of University Association for Science and Technology in Shaanxi, China (Grant No.20170402).

## Author contributions

Ke Wang, Yixuan Wang and Zebing Hu contributed equally to the study design, experimental work and data analysis. Fei Shi, Shu Zhang and Ge Zhang supervised the project. Ke Wang wrote the manuscript. Fei Shi, Shu Zhang reviewed the manuscript. Xinsheng Cao, Lijun Zhang, Gaozhi Li and Yingjun Tan contributed to unloading model establishment and sample collection of aimals experiments. Bone targeted materials were synthesized and provided by Lei Dang and Ge Zhang.

## Conflict of interest

The authors have declared that no competing interest exists.

